# Phylogeny predicts tolerance in aquatic animals for only a minority of chemicals

**DOI:** 10.1101/2024.02.20.577278

**Authors:** Alice L. Coleman, Suzanne Edmands

## Abstract

There are substantial gaps in our empirical knowledge of the effects of chemical exposure on aquatic life that are unlikely to be filled by traditional laboratory toxicity testing alone. One possible alternative of generating new toxicity data is cross-species extrapolation (CSE), a statistical approach in which existing data are used to predict the effect of a chemical on untested species. Some CSE models use relatedness as a predictor of chemical sensitivity, but relatively little is known about how strongly shared evolutionary history influences sensitivity across all chemicals. To address this question, we conducted a survey of phylogenetic signal in the toxicity data from aquatic animal species for a large set of chemicals using a phylogeny inferred from taxonomy. Strong phylogenetic signal was present in just six of thirty-two toxicity datasets, and there were no clear shared properties among those datasets with strong signal. Strong signal was rare even among chemicals specifically developed to target insects, meaning that these chemicals may be equally lethal to non-target taxa, including chordates. When signal was strong, distinct patterns of sensitivity were evident in the data, which may be informative when assembling toxicity datasets for regulatory use. Although strong signal does not appear to manifest in aquatic toxicity data for most chemicals, we encourage additional phylogenetic evaluations of toxicity data in order to guide the selection of CSE tools and as a means to explore the patterns of chemical sensitivity across the broad diversity of life.

## Introduction

Pollution represents one of the greatest threats to global biodiversity (Novacek and Cleland 2001). The biological consequences of pollution exposure are numerous, including reduced fecundity, compromised immune systems, developmental abnormalities and outright mortality (McKinlay et al. 2008), all of which threaten the health and functioning of organisms, populations and ecosystems at large. In recognition of this danger, regulatory bodies like the United States Environmental Protection Agency (EPA) manage pollution in part by setting numeric criteria that are intended to protect life by limiting the accumulation of hazardous concentrations of chemicals in the environment. The development of such criteria depends on empirical data that describe the concentrations of chemicals that inflict adverse effects on organisms, however, most chemicals do not have robust toxicity datasets (Coleman and Edmands 2022; Wheeler et al. 2002; Dowse et al. 2013).

Most toxicity data are derived from single-species toxicity tests that measure the dose of a substance that induces adverse effects in a test population over a certain period. These tests are classified as either acute (short-term) or chronic (long-term), with chronic tests typically performed less frequently because of the high costs associated with long-term testing. In theory, the species evaluated in such tests should be representative of the biological community a criterion is intended to protect, but in practice, most testing is performed using a select group of model species (Seegert, Fava, and Cumbia 1985; Buchwalter, Clements, and Luoma 2017; Anderson and Phillips 2016). As a result, our existing toxicity database does not necessarily reflect the broad diversity of life nor the entire spectrum of sensitivity to any given chemical. The logical means of addressing these data gaps would be to simply increase testing, but the sheer number of possible species-chemical combinations means that expanding laboratory efforts is not a viable solution. As a result, there is a demand for other tools that can reliably estimate how sensitive a species is to any given chemical. At present, computational methods that extrapolate existing toxicity data to untested species (i.e cross-species extrapolation) represent the most promising alternative to traditional laboratory testing (LaLone et al. 2021; van den Berg et al. 2021).

There are four main types of cross-species extrapolation (CSE) models: interspecies correlation, trait, genomic and relatedness-based (van den Berg et al. 2021). Here, we are most interested in relatedness-based models, which operate under the assumption that evolutionary relationships can explain the variation in chemical sensitivity across taxa. These models generate new toxicity data for species by using metrics of relatedness as predictors of sensitivity. One such metric used in relatedness-based CSE is phylogenetic signal, which is a measure of the statistical dependence among species trait values that arises from their phylogenetic relationships (Revell, Harmon, and Collar 2008; van den Berg et al. 2021). When phylogenetic signal is high, close relatives on a phylogeny will tend to exhibit very similar trait values while distantly related species will not. When signal is low, trait values will tend to be randomly distributed across a phylogeny and distantly related taxa may resemble each other more than close relatives (Kamilar and Cooper 2013; Kamilar and Muldoon 2010). Chemical sensitivity is regarded as a possible candidate to exhibit strong phylogenetic signal (Hylton et al. 2018).

Sensitivity to a chemical is a complex phenotype determined by a network of interacting morphological, physiological and behavioral traits that regulate the uptake, metabolism and excretion of pollutants by an organism. Many traits linked to sensitivity such as body size and metabolic rate are known to exhibit phylogenetic signal, suggesting that we are likely to find some amount of signal in chemical sensitivity data (Hylton et al. 2018; Kamilar and Cooper 2013). To date, only a small number of studies have evaluated toxicity data for phylogenetic signal, some of which identified meaningful associations between shared evolutionary history and species sensitivity (Hylton et al. 2018; Chiari et al. 2015; Guénard et al. 2011; Hammond et al. 2012; Buchwalter et al. 2008; Moore et al. 2020). The scope of these studies has been relatively narrow, with emphasis placed on data from either a single chemical or taxonomic group. These limitations mean that we do not have a full understanding of either the variation in chemical sensitivity across the diversity of life or the abundance of strong phylogenetic signal in toxicity data. As a result, the practicality of using relatedness-based cross-species extrapolation to fill data gaps on a large scale remains in question.

Improved knowledge of phylogenetic signal in toxicity data may have other uses aside from cross-species extrapolation. One option would be to test hypotheses that explain why phylogenetic signal manifests more strongly in some datasets than others. To this end, we propose three hypotheses based on qualitative properties of chemicals and toxicity data. First, we predict that strong phylogenetic signal will be more common among synthetic chemicals than naturally occurring chemicals. Synthetic chemicals are relative newcomers in the environment, so potentially only a subset of lineages will have had enough exposure for selection to favor increased tolerance. Second, we predict that strong phylogenetic signal will be more common in the data from chemicals with specific toxic modes of action (MOAs) than those that are nonspecific. This is because chemicals specific MOAs interfere with biological processes by precisely binding to a particular site or molecule, so an adaptive phenotype could in theory arise after only a small number of molecular changes (Gupta 2018; Whitehead et al. 2017). Chemicals with nonspecific MOAs tend to induce generic stress responses, so the corresponding adaptive phenotypes may be complex and thus take much longer to develop (Whitehead et al. 2017). Lastly, we predict that strong phylogenetic signal will be more common in chronic toxicity datasets than acute because rapid responses to acute stress are often generalized and evolutionarily conserved (Kültz 2020). In contrast, responses to chronic stress have been found to be more chemical- and lineage-specific (McRae et al. 2022; Kovalchuk et al. 2007). Testing hypotheses such as these may provide insight into whether qualitative properties are viable predictors of strong phylogenetic signal, which would be useful for identifying the datasets to which relatedness-based models can be applied.

In this study we expand upon previous phylogenetic analyses in ecotoxicology by considering toxicity data from both a large number of chemical pollutants and a biologically diverse set of species. Using published laboratory toxicity data and a phylogeny inferred from taxonomy, we quantified the phylogenetic signal present in these data and explored patterns of sensitivity across taxa. We also investigated whether life stage, exposure type, chemical origin, or mode of action (MOA) affected signal and evaluated the influences of experimental temperature and pH on chemical sensitivity using phylogenetic methods. The application of CSE methods to environmental risk assessment and regulation has been a recent focus in aquatic toxicology (Raimondo, Jackson, and Barron 2010; Coleman and Edmands 2022; Schlekat et al. 2010; Lewis and Thursby 2018), so here our analysis specifically dealt with data similar to those used in water quality criteria development.

## Methods

### Data Collection

Toxicity data from aquatic species were collected for twenty-four chemicals (Table 1) that span a range of origins, classes and modes of action (MOA). The MOA of each chemical was obtained and labeled as either generic or precise using the MOAtox database from Barron et al. (2015). Acute toxicity data were then obtained from the EPA’s ECOTOXicology Knowledgebase (ECOTOX; Olker et al. 2022) using these search parameters:

1. Chemical Abstracts Service (CAS) Registry Number
2. Endpoint: Median Lethal Concentration (LC50)
3. Kingdom: Animals
4. Test Location: Lab
5. Exposure Media: Freshwater or Saltwater
6. Duration: 4 days

**Table 1.** Properties of chemicals evaluated in analyses.

| Chemical | CAS RN | Origin | Class | Mode of Action |
| --- | --- | --- | --- | --- |
| Ammonia | 7664-41-7 | Natural | Inorganic | Osmoregulatory impairment<sup>a</sup> |
| Cadmium | 7440-43-9 | Natural | Metal | Metallic iono/osmoregulatory impairment<sup>a</sup> |
| Chlorine | 7782-50-5 | Natural | Halogen | Osmoregulatory impairment<sup>a</sup> |
| Copper | 7440-50-8 | Natural | Metal | Metallic iono/osmoregulatory impairment<sup>a</sup> |
| Mercury | 7439-97-6 | Natural | Metal | Metallic iono/osmoregulatory impairment<sup>a</sup> |
| Nickel | 7440-02-0 | Natural | Metal | Metallic iono/osmoregulatory impairment<sup>a</sup> |
| Phenol | 108-95-2 | Natural | Phenol | Polar narcosis<sup>a</sup> |
| Toluene | 108-88-3 | Natural | Aromatic Hydrocarbon | Nonpolar narcosis<sup>a</sup> |
| Zinc | 7440-66-6 | Natural | Metal | Metallic iono/osmoregulatory impairment<sup>a</sup> |
| 4-Nitrophenol | 100-02-7 | Synthetic | Nitrophenol | Polar narcosis<sup>a</sup> |
| Atrazine | 1912-24-9 | Synthetic | Triazine | Narcosis<sup>a</sup> |
| Chlorpyrifos | 2921-88-2 | Synthetic | Organophosphate | AChE Inhibition<sup>b</sup> |
| DDT | 50-29-3 | Synthetic | Organochlorine | Neurotoxicity<sup>b</sup> |
| Diazinon | 333-41-5 | Synthetic | Organophosphate | AChE inhibition<sup>b</sup> |
| Dieldrin | 60-57-1 | Synthetic | Organochlorine | Neurotoxicity<sup>b</sup> |
| Endosulfan | 115-29-7 | Synthetic | Organochlorine | Neurotoxicity<sup>b</sup> |
| Endrin | 72-20-8 | Synthetic | Organochlorine | Neurotoxicity<sup>b</sup> |
| Glyphosate | 1071-83-6 | Synthetic | Phosphonate | Nonpolar narcosis<sup>a</sup> |
| Guthion | 86-50-0 | Synthetic | Organophosphate | AChE inhibition<sup>b</sup> |
| Lindane | 58-89-9 | Synthetic | Organochlorine | Neurotoxicity<sup>b</sup> |
| Malathion | 121-75-5 | Synthetic | Organophosphate | AChE inhibition<sup>b</sup> |
| Parathion | 56-38-2 | Synthetic | Organophosphate | AChE inhibition<sup>b</sup> |
| PCP | 87-86-5 | Synthetic | Organochlorine | Electron transport inhibition<sup>b</sup> |
| TBTO | 56-35-9 | Synthetic | Organotin | Electron transport inhibition<sup>b</sup> |

Abbreviations: CAS RN = Chemical Abstracts Service Registry Number, DDT = 1,1'-(2,2,2-
Trichloroethane-1,1-diyl)bis(4-chlorobenzene), PCP = Pentachlorophenol, TBTO = Tributyltin
oxide, AChE = Acetylcholinesterase
<sup>a</sup> Generic mode of action

These restrictions ensured that we obtained comparable data for different species tested with the same chemical and were specifically designed to approximate how the EPA collects data for water quality criteria development (Stephan et al. 1985). Acute test duration was set to 96 hours following Chiari et al. (2015) and Hylton et al. (2018). Toxicity values without a precise species name were omitted. When a single species contributed more than one toxicity value to a particular dataset, we calculated the species value as the geometric mean of all its datapoints.

Chronic toxicity data were similarly mined from ECOTOX for eight chemicals (Table 2) using the following parameters:

1. CAS Registry Number
2. Endpoint: No Observed Effect Concentration (NOEC)
3. Kingdom: Animals
4. Test Location: Lab
5. Exposure Media: Freshwater or Saltwater
6. Duration: 14 – 30 days

**Table 2.** Phylogenetic signal results for all dataset variations.

| Chemical | Exposure | Dataset Type |  |  |  |
| --- | --- | --- | --- | --- | --- |
|  |  | Complete | Subadult | Temperature | pH |
| | | $\lambda$ | $\lambda$ | $\lambda$ | $\lambda$ |
| Ammonia | Acute | 6.6E-05 | 6.7E-05 | 0.0 | 0.0 |
| Cadmium | Acute | 6.6E-05 | 0.41 | 0.0 | 0.0 |
| Cadmium | Chronic | **0.98** | **1.0** | 0.0 | 0.0 |
| Chlorine | Acute | **0.923** | **0.68** | **0.92** | **0.77** |
| Copper | Acute | 0.0023 | 7.2E-04 | 0.013 | 0.12 |
| Copper | Chronic | **1.0** | 6.7E-05 | 0.0 | 0.0 |
| Mercury | Acute | 6.6E-05 | 6.7E-05 | 0.0 | 0.0 |
| Nickel | Acute | 6.6E-05 | 0.22 | 0.049 | 0.0 |
| Phenol | Acute | 0.23 | **0.60** | 0.22 | 0.12 |
| Toluene | Acute | 6.7E-05 | 6.7E-05 | 0.0 | 0.0 |
| Zinc | Acute | 6.6E-05 | 6.7E-05 | 0.0 | 0.0 |
| 4-Nitrophenol | Acute | 6.7E-05 | **0.94** | 0.0 | 0.0 |
| Atrazine | Acute | 0.0052 | 6.6E-05 | 0.0 | 0.0 |
| Atrazine | Chronic | 0.018 | **0.75** | **0.99** | 1.0 |
| Chlorpyrifos | Acute | 6.6E-05 | 6.6E-05 | 0.0 | 0.0 |
| Chlorpyrifos | Chronic | **1.0** | 0.28 | **1.0** | 0.0 |
| DDT | Acute | 6.6E-05 | **0.54** | 0.0 | 0.0 |
| Diazinon | Acute | 0.025 | 0.039 | 0.023 | 0.40 |
| Diazinon | Chronic | 6.6E-05 | 4.5E-05 | 0.0 | 1.0 |
| Dieldrin | Acute | **0.87** | 6.6E-05 | 0.16 | 0.21 |
| Endosulfan | Acute | 6.6E-05 | **0.94** | 0.0 | 0.0 |
| Endrin | Acute | 6.6E-05 | 5.6E-05 | 0.0 | 0.0 |
| Glyphosate | Acute | 6.6E-05 | **0.61** | 0.0 | 0.0 |
| Glyphosate | Chronic | 6.6E-05 | 6.6E-05 | 0.0 | 0.0 |
| Guthion | Acute | **0.79** | **0.69** | **0.55** | **0.57** |
| Lindane | Acute | 6.6E-05 | **0.99** | 0.0 | 0.0 |
| Malathion | Acute | 0.23 | 0.35 | 0.18 | 0.30 |
| Malathion | Chronic | 6.6E-05 | 6.7E-05 | 0.11 | 0.171 |
| Parathion | Acute | 0.20 | **0.65** | 0.15 | **0.67** |
| PCP | Acute | 0.067 | 0.24 | 0.027 | 0.043 |
| PCP | Chronic | 6.7E-05 | 6.7E-05 | 0.0 | 0.0 |
| TBTO | Acute | 6.6E-05 | 6.7E-05 | 0.0 | 0.0 |

Abbreviations: DDT = 1,1'-(2,2,2-Trichloroethane-1,1-diyl)bis(4-chlorobenzene), PCP =
Pentachlorophenol, TBTO = Tributyltin oxide
Values in **boldface** indicate strong phylogenetic signal ( $\lambda > 0.5$ )

The remaining 16 chemicals in our initial set of 24 were excluded because of low availability of chronic data in ECOTOX. As there is no universally agreed upon period that constitutes a chronic toxicity test, we allowed for a broad range of durations to maximize the amount of usable data.

Test organism life stage, experimental temperature and pH for each toxicity value were also obtained from ECOTOX when possible because these variables are known to modify the toxicity of many chemicals to aquatic species (Wang, Meador, and Leung 2016; Cairns, Heath, and Parker 1975; Cadmus et al. 2020; Alonso, De Lange, and Peeters 2010). Life stage was standardized to either “subadult”, “adult” or “NR” (not reported) based on the information provided by ECOTOX. Experimental temperature and pH were summarized as geometric means for species with multiple values for either variable.

### Phylogenetic Analyses

We used the National Center for Biotechnology Information’s (NCBI) Taxonomy database via the phyloT generator (Letunica 2022) to produce a single comprehensive phylogenetic tree for all of the species that appeared in the toxicity datasets we assembled. Species without a matching entry in the NCBI database were excluded from the tree. We imported the tree generated by phyloT into R using the package ape (Paradis and Schliep 2019), where we then computed its branch lengths using Grafen’s transformation (ρ = 1; Grafen and Hamilton 1989) and annotated its tips with the toxicity data by matching to species names.

Phylogenetic signal was estimated as Pagel’s λ (Pagel 1998). The values of λ range between zero and one, where a value close to zero indicates that the trait is phylogenetically independent and a λ near one indicates that the distribution of the trait is consistent with the Brownian Motion model, wherein closely related lineages are more similar to each other than those that are distantly related (Münkemüller et al. 2012). Following Hylton et al. (2018), we considered λ values of 0.5 or greater to be evidence of strong phylogenetic signal. Two initial calculations of λ were performed for each toxicity dataset using the function phylosig from the R package phytools (Revell 2012). The first calculation was performed for the complete dataset while the second was performed for the subset of toxicity data that were derived from subadult test organisms. When strong signal was identified, we plotted the toxicity data alongside the phylogeny we assembled (Fig. 1-6, S1-S3) and assessed the visual patterns in the data. We used the second calculation of λ to determine whether the strength of signal or patterns of sensitivity differed in organisms tested at early (subadult) versus various (subadult, adult and NR) life stages.

**Fig. 1.**
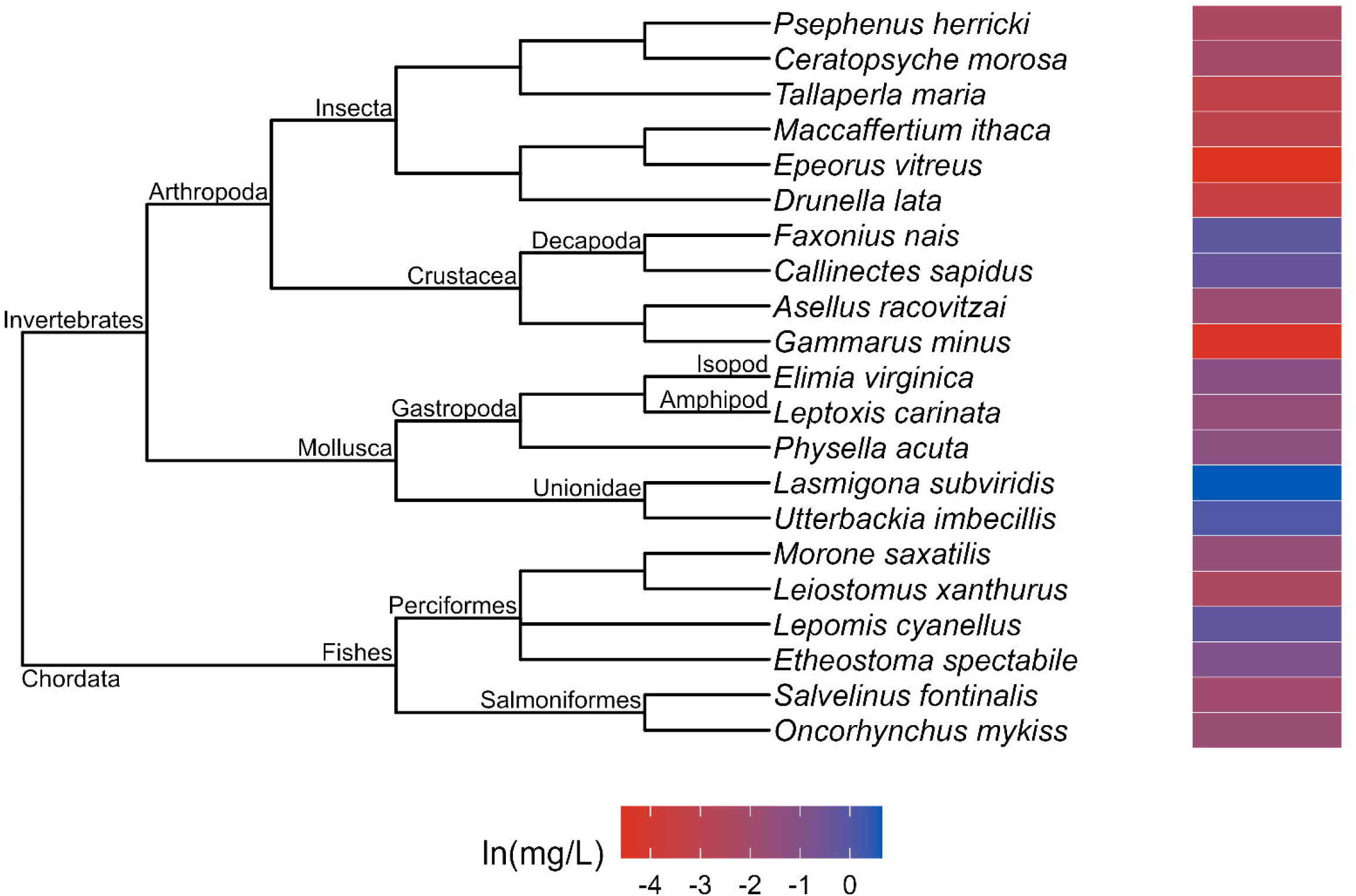
Phylogenetic tree and toxicity data heatmap for the acute chlorine dataset (λ = 0.53). The colored bar next to each species represents its relative sensitivity to the chemical. A red bar indicates a high degree of sensitivity (i.e. small amount of chemical causes toxic effect), while a blue bar indicates low sensitivity (i.e. large amount of chemical causes toxic effect)

The effects of temperature and pH on chemical sensitivity were evaluated using linear phylogenetic generalized least squares (PGLS; Grafen and Hamilton 1989) models. A form of regression analysis, PGLS models are used to test the association between variables while statistically controlling for the effects of phylogenetic signal in the data (Symonds and Blomberg 2014). Using the package caper (Orme et al. 2018; Grafen and Hamilton 1989), we set up models for the subsets of each toxicity dataset that had temperature or pH information available, using toxicity (LC50 or NOEC) as the dependent variable and the environmental information (°C or pH) as the predictor variable. These models also included a measurement of λ in each data subset, which in PGLS is calculated on model residuals using maximum likelihood estimation (Revell 2010; Symonds and Blomberg 2014; Pagel 1999). Temperature and pH were modeled separately because of most toxicity values did not have information for both variables available.

## Results

### Data composition

We collected a cumulative total of 5,800 datapoints to create 32 toxicity datasets (24 acute, 8 chronic) for 24 chemicals (9 natural, 15 synthetic; 12 generic MOAs, 12 precise MOAs). Together, these datasets included values from 512 unique species from ten phyla (Annelida, Arthropoda, Bryozoa, Chordata, Cnidaria, Echinodermata, Mollusca, Nematoda, Platyhelminthes and Rotifera). The sizes of the toxicity datasets varied considerably, with the largest (acute DDT) composed of 368 datapoints from 99 species while the smallest (acute 4-nitrophenol) was made up of just 42 datapoints from 14 species. In general, acute toxicity datasets were larger than chronic and synthetic chemical datasets were larger those of the naturally-occurring chemicals (Table S1). Notably, the chronic toxicity dataset for atrazine was larger (*n* = 350 datapoints) than its acute counterpart (*n* = 109 datapoints) but contained fewer species (chronic *n* = 43 species, acute *n* = 53 species). Dataset sample sizes decreased substantially when filtered by availability any one of life stage, temperature or pH information (Table S1).

### Phylogenetic signal

Strong phylogenetic signal (λ ≥ 0.5) was identified in six of the 32 toxicity datasets (acute chlorine, acute dieldrin, acute guthion, chronic cadmium, chronic chlorpyrifos and chronic copper) when we did not consider any of life stage, experimental temperature or pH (Table 2). Convergence on the lower bound of λ was common, as 17 datasets returned a signal value at or near 6.6E-05. Fisher’s exact tests indicated that there was no difference in frequencies of strong phylogenetic signal between toxicity datasets from chemicals of different origins (natural vs. synthetic; *p* = 1), classes (see “Class” column in Table 1; *p* = 0.86) or modes of action (generic vs. precise; *p* = 1). Exposure type (acute vs. chronic; *p* = 0.085) similarly did not affect the frequency of strong phylogenetic signal.

Strong phylogenetic signal was more common in the data derived from subadult test organisms, with eleven datasets returning a lambda value greater than 0.5 (Table 2). However, the prevalence of strong signal in these datasets was not significantly different from the frequency in the complete datasets (Fisher’s exact test; *p* = 1). There were three instances of overlapping strong phylogenetic signal between the complete and subadult versions of a dataset (acute chlorine, acute guthion and chronic chlorpyrifos). Additionally, there were three instances where strong signal was present in a complete dataset but not in its respective subadult dataset, and eight cases of the reverse situation wherein subadult datasets showed strong signal but their corresponding complete datasets did not. Notably, several of the subadult datasets from this third scenario had very small species sample sizes (*n* ≤ 10; Table S1)

In contrast to the subadult results, strong phylogenetic signal was less common in both the temperature (3/32) and pH (3/32) data subsets than in the complete datasets (6/32). Here, there were two cases (acute chlorine and acute guthion) of overlapping signal between the complete and temperature/pH datasets. Lambda values were largely consistent across all dataset variations in those instances. We also identified one case of strong signal in a temperature data subset (chronic chlorpyrifos) and one in a pH data subset (acute parathion) that were not present in the original datasets.

### Patterns of sensitivity

The patterns of sensitivity in the acute toxicity datasets with strong phylogenetic signal varied across chemicals. For example, the acute data for chlorine (λ = 0.93; Fig. 1) suggests that insects are the most sensitive group on the phylogeny while fishes, crustaceans and molluscs are more tolerant. Within the crustaceans, the acute chlorine tolerance of decapods appears to exceed that of both isopods and amphipods while in the molluscs, the Unionidae family of freshwater mussels appear more resistant than the gastropods. The acute dieldrin data (λ = 0.87; Fig. 2) shows a different pattern, wherein fishes, outside of the Cypriniformes, appear to be the most sensitive taxa while many frogs (order Anura) and some crustaceans exhibit relatively high tolerance. In the acute guthion data (λ = 0.79; Fig. 3) the arthropods, and in particular many crustacean species, are the most vulnerable to the organophosphate pesticide while the amphibians and many species of fish from the orders Cypriniformes and Siluriformes fall on the opposite end of the spectrum of sensitivity.

**Fig. 2.**
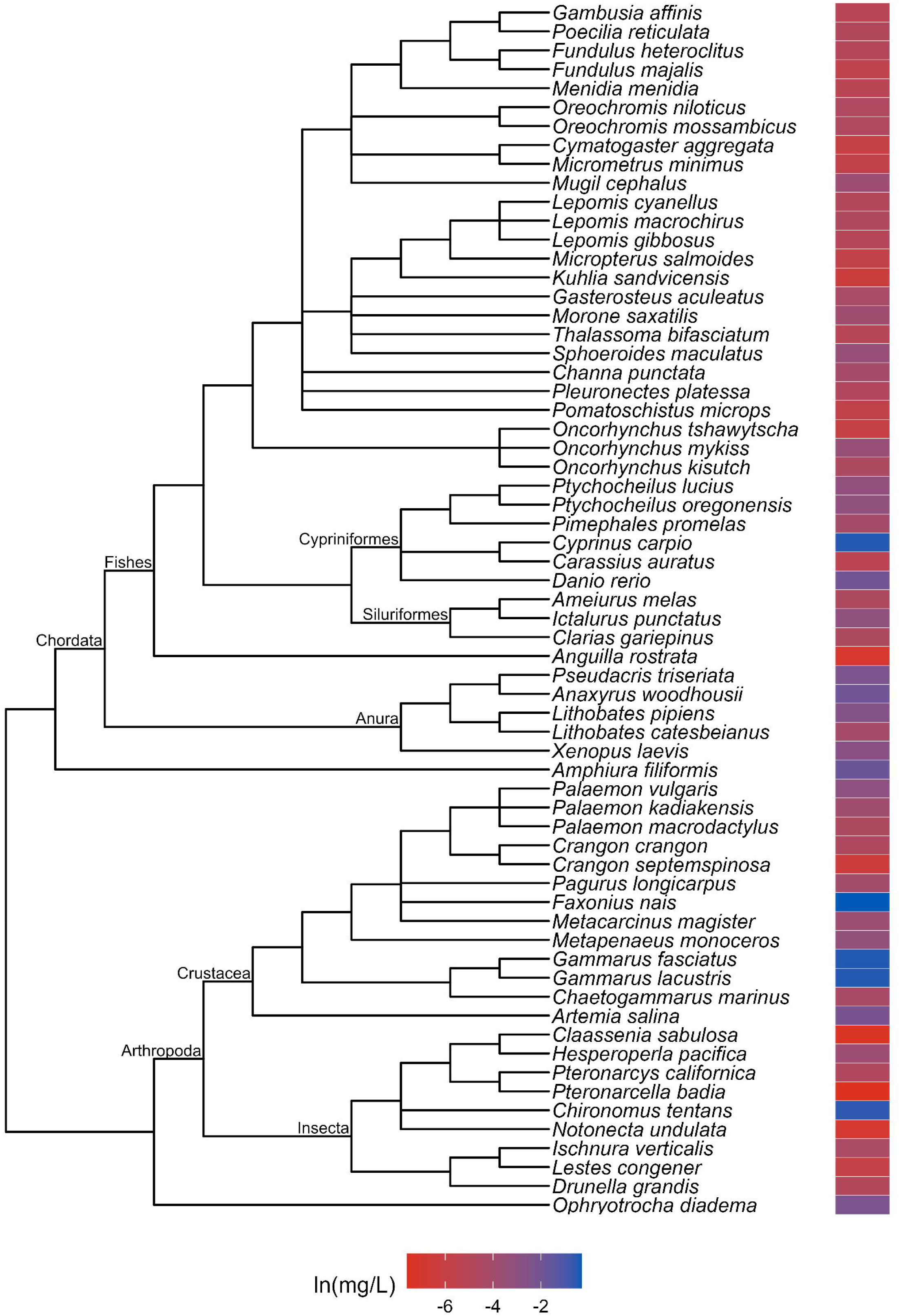
Phylogenetic tree and toxicity data heatmap for the acute dieldrin dataset (λ = 0.931). The colored bar next to each species represents its relative sensitivity to the chemical. A red bar indicates a high degree of sensitivity (i.e. small amount of chemical causes toxic effect), while a blue bar indicates low sensitivity (i.e. large amount of chemical causes toxic effect)

**Fig. 3.**
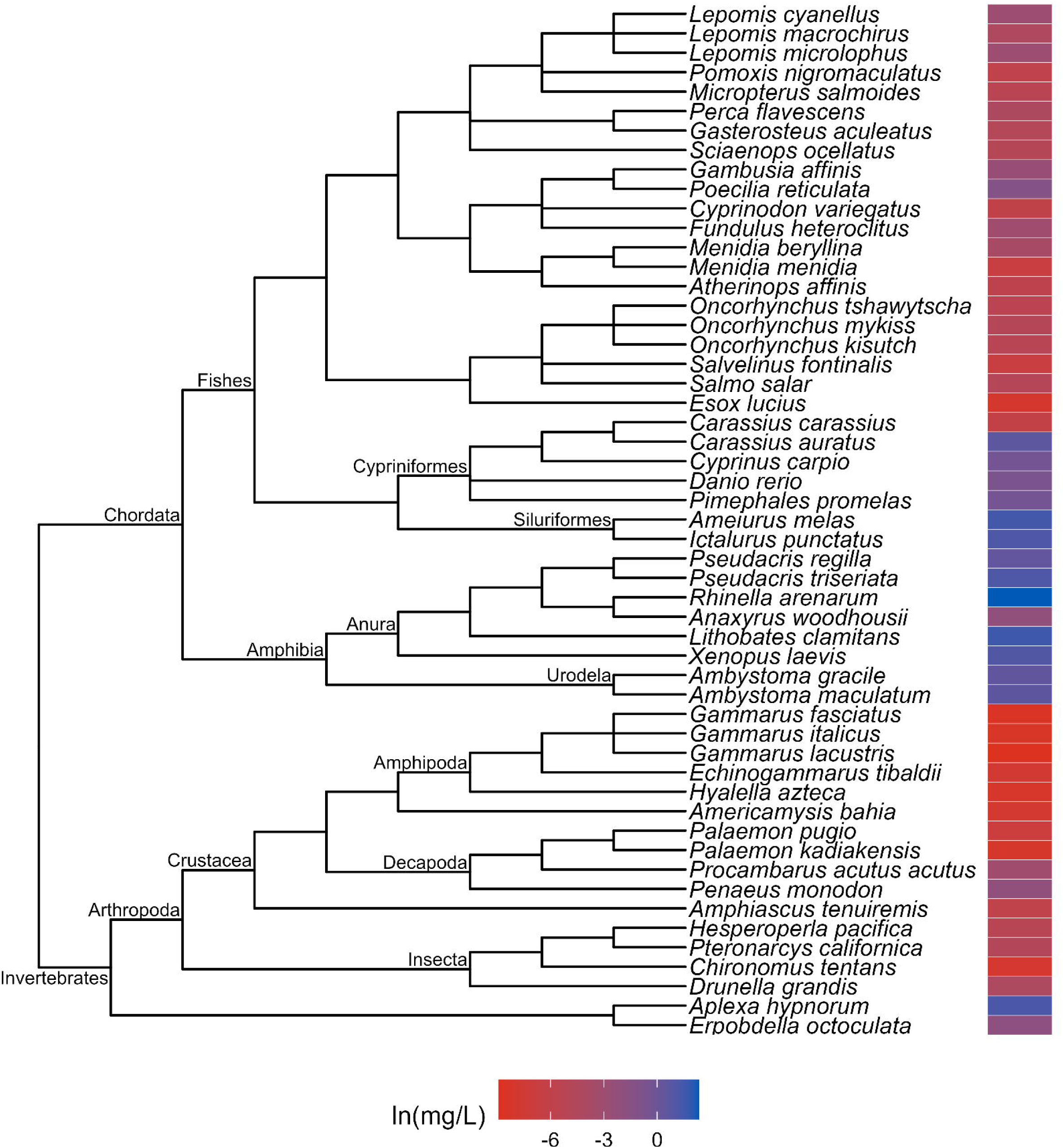
Phylogenetic tree and toxicity data heatmap for the acute guthion dataset (λ = 1). The colored bar next to each species represents its relative sensitivity to the chemical. A red bar indicates a high degree of sensitivity (i.e. small amount of chemical causes toxic effect), while a blue bar indicates low sensitivity (i.e. large amount of chemical causes toxic effect)

Similar pattern variation was present in the chronic toxicity data. The chronic cadmium plot (λ = 0.98; Fig. 4) indicates that crustaceans are highly sensitive to the metal while molluscs and fishes are more tolerant. Fishes and other chordates also exhibit high tolerance in the chronic chlorpyrifos plot (λ = 1.0; Fig. 5), while rotifers and arthropods appear to be the most sensitive taxa. The majority of species featured in the chronic copper dataset (λ = 1.0; Fig. 6) seem to experience toxic effects when exposed to low levels of copper, with exceptionally high tolerance evident in just one species of frog (*Pelophylax ridibundus*) and a few cases of moderate tolerance scattered across different taxa.

**Fig. 4.**
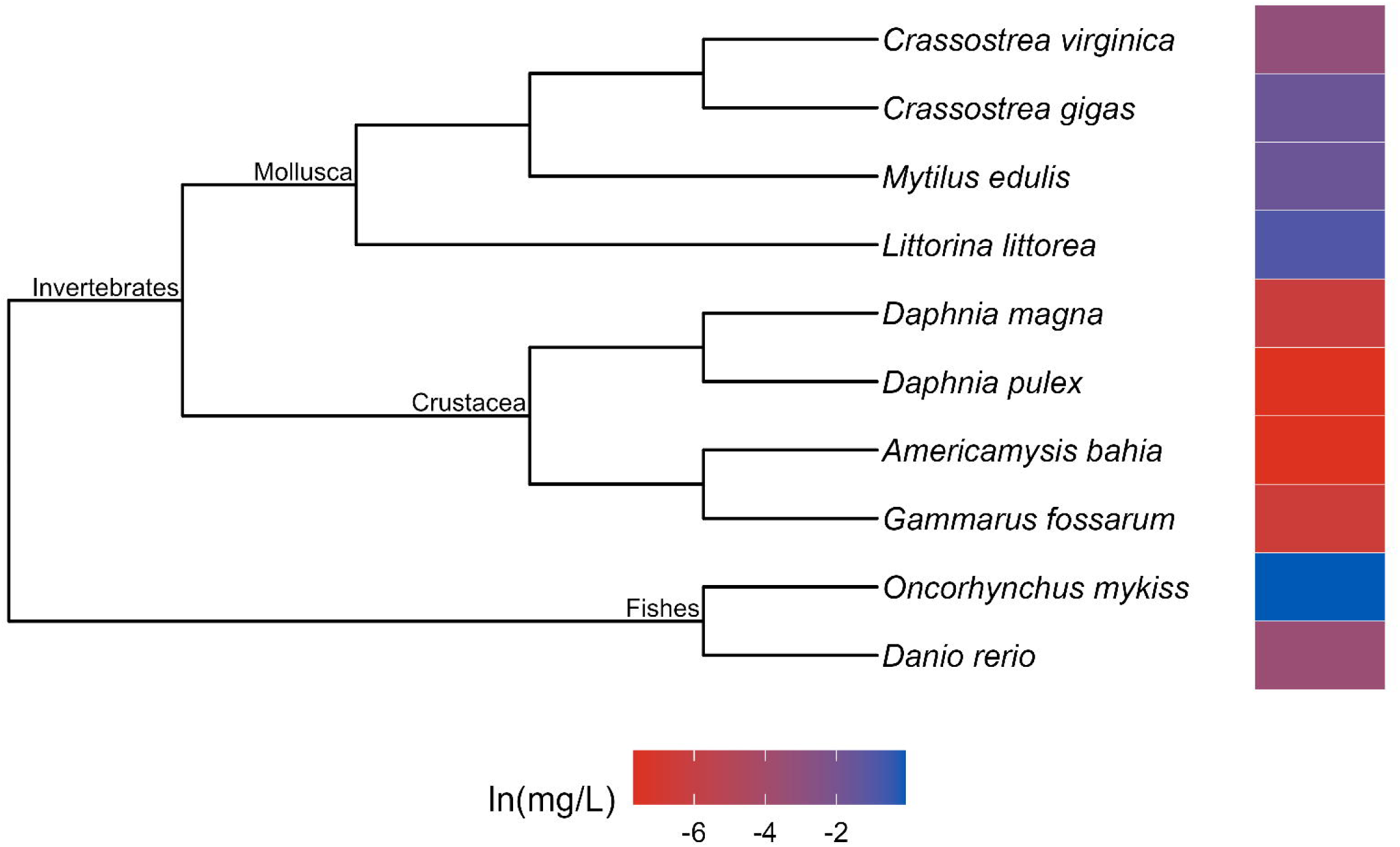
Phylogenetic tree and toxicity data heatmap for the chronic cadmium dataset (λ = 0.995). The colored bar next to each species represents its relative sensitivity to the chemical. A red bar indicates a high degree of sensitivity (i.e. small amount of chemical causes toxic effect), while a blue bar indicates low sensitivity (i.e. large amount of chemical causes toxic effect)

**Fig. 5.**
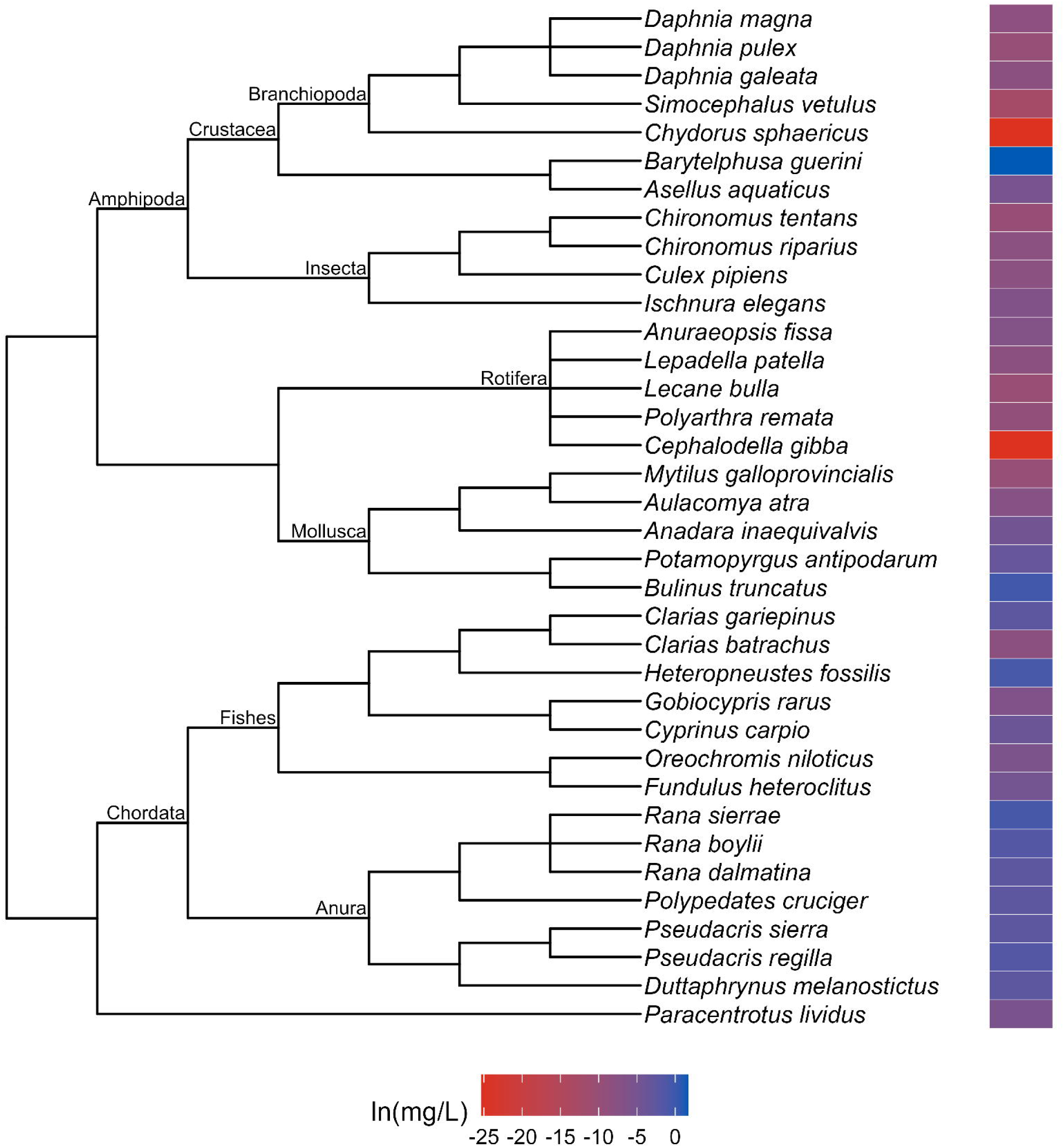
Phylogenetic tree and toxicity data heatmap for the chronic chlorpyrifos dataset (λ = 1). The colored bar next to each species represents its relative sensitivity to the chemical. A red bar indicates a high degree of sensitivity (i.e. small amount of chemical causes toxic effect), while a blue bar indicates low sensitivity (i.e. large amount of chemical causes toxic effect)

**Fig. 6.** Combined phylogenetic tree and toxicity data heatmap for the chronic copper dataset (λ = 1). The colored bar next to each species represents its relative sensitivity to the chemical. A red bar indicates a high degree of sensitivity (i.e. small amount of chemical causes toxic effect), while a blue bar indicates low sensitivity (i.e. large amount of chemical causes toxic effect)

Additionally, sensitivity patterns differed minimally between the complete and subadult variations of datasets that exhibited strong phylogenetic signal in both forms (acute chlorine, acute guthion and chronic cadmium; Fig. S1-S3).

### PGLS

Experimental temperature and pH were significant predictors of toxicity in PGLS models for four and three datasets each (Table 3). The adjusted R^2^ values of the significant models ranged between 0.037 and 0.7 for temperature and between 0.14 and 0.37 for pH (Table 3). Experimental temperature was positively associated with tolerance (i.e. LC50/NOEC increases when temperature increases) in the acute dieldrin and acute malathion datasets and negatively associated with tolerance (i.e. LC50/NOEC decreases when temperature increases) in the acute cadmium and acute phenol datasets. Experimental pH was positively associated with tolerance in the acute chlorine data and negatively associated with tolerance in the acute ammonia, chronic diazinon and chronic pentachlorophenol datasets.

**Table 3.** Results of PGLS regressions for temperature and pH with toxicity data.

| Chemical | Exposure | Temperature |  |  | pH |  |  |
| --- | --- | --- | --- | --- | --- | --- | --- |
|  |  | Coefficient | *p* | A-R<sup>2</sup> | Coefficient | *p* | A-R<sup>2</sup> |
| Ammonia | Acute | 6.4 | 0.056 | 0.073 | **-110** | **0.012** | **0.14** |
| Cadmium | Acute | **-2.2** | **0.021** | **0.070** | -6.6 | 0.80 | -0.023 |
| Cadmium | Chronic | 10 | 0.38 | -0.013 | 196 | 0.50 | -0.12 |
| Chlorine | Acute | 0.0059 | 0.70 | -0.044 | **0.75** | **0.0029** | **0.37** |
| Copper | Acute | 0.14 | 0.53 | -0.0090 | 0.51 | 0.21 | 0.013 |
| Copper | Chronic | 0.22 | 0.78 | -0.048 | 4.3 | 0.84 | -0.050 |
| Mercury | Acute | -0.0036 | 0.64 | -0.037 | -0.024 | 0.60 | -0.054 |
| Nickel | Acute | 0.48 | 0.67 | -0.050 | -113 | 0.08 | 0.18 |
| Phenol | Acute | **-4.6** | **0.049** | **0.054** | 37 | 0.46 | -0.010 |
| Toluene | Acute | 16 | 0.17 | 0.050 | 45 | 0.91 | -0.090 |
| Zinc | Acute | 0.55 | 0.20 | 0.014 | -1.8 | 0.89 | -0.033 |
| 4-Nitrophenol | Acute | 0.058 | 0.89 | -0.082 | 1.2 | 0.92 | -0.12 |
| Atrazine | Acute | 0.72 | 0.79 | -0.019 | -5.0 | 0.57 | -0.017 |
| Atrazine | Chronic | -0.74 | 0.23 | 0.014 | -1.6 | 0.69 | -0.038 |
| Chlorpyrifos | Acute | -0.04 | 0.60 | -0.008 | 0.44 | 0.72 | -0.012 |
| Chlorpyrifos | Chronic | -0.0052 | 0.78 | -0.037 | -0.28 | 0.57 | -0.041 |
| DDT | Acute | 0.10 | 0.54 | -0.0060 | 0.79 | 0.77 | -0.013 |
| Diazinon | Acute | -0.53 | 0.12 | 0.018 | 0.40 | 0.68 | -0.015 |
| Diazinon | Chronic | 0.41 | 0.058 | 0.39 | **-1.8** | **0.050** | **0.57** |
| Dieldrin | Acute | **0.0099** | **0.043** | **0.053** | -0.085 | 0.064 | 0.045 |
| Endosulfan | Acute | -0.029 | 0.78 | -0.011 | -0.41 | 0.53 | -0.0090 |
| Endrin | Acute | 0.011 | 0.25 | 0.0052 | -0.071 | 0.52 | -0.011 |
| Glyphosate | Acute | -11 | 0.82 | -0.035 | 3.7 | 0.94 | -0.062 |
| Glyphosate | Chronic | 1.4 | 0.38 | -0.014 | -2.1 | 0.48 | -0.087 |
| Guthion | Acute | -0.034 | 0.45 | -0.0090 | 0.82 | 0.31 | 0.0016 |
| Lindane | Acute | 0.075 | 0.65 | -0.0090 | 1.3 | 0.57 | -0.0080 |
| Malathion | Acute | **0.72** | **0.016** | **0.037** | -0.84 | 0.85 | -0.0090 |
| Malathion | Chronic | -0.033 | 0.76 | -0.089 | -4.9 | 0.12 | 0.37 |
| Parathion | Acute | 0.14 | 0.24 | 0.0084 | 0.42 | 0.52 | -0.017 |
| PCP | Acute | -0.72 | 0.068 | 0.027 | 2.7 | 0.59 | -0.090 |
| PCP | Chronic | 1.6 | 0.48 | -0.028 | **-41** | **0.0051** | **0.33** |
| TBTO | Acute | -4.0 | 0.39 | -0.0080 | -26 | 0.77 | -0.034 |

Abbreviations: A-R<sup>2</sup>= Adjusted R<sup>2</sup>, DDT = 1,1'-(2,2,2-Trichloroethane-1,1-diyl)bis(4-
chlorobenzene), PCP = Pentachlorophenol, TBTO = Tributyltin oxide
Values in **boldface** indicate a significant ( $p \leq 0.05$ ) positive relationship
Values in **boldface italics** indicate a significant ( $p \leq 0.05$ ) negative relationship

## Discussion

Our results indicate that strong phylogenetic signal is rare in toxicity data from aquatic animals and does not appear to be biased by organism life stage, exposure type, chemical origin, class or mode of action. We also found evidence of significant effects of experimental temperature and pH on chemical tolerance in a few datasets, although there were not consistent trends in how either factor affected toxicity across chemicals. High variation in phylogenetic signal magnitude across different chemicals is consistent with the results of the similarly-sized analysis by Hylton et al. (2018), which found strong signal in just 10 of the 42 toxicity datasets they examined. In general, most of the previous studies (Hylton et al. 2018; Chiari et al. 2015; Guénard et al. 2011; Hammond et al. 2012; Buchwalter et al. 2008) that identified strong phylogenetic signal appear to have less taxonomic breadth in their data than what we used in our work, suggesting that phylogenetic signal might manifest more strongly at lower taxonomic levels (Carew, Miller, and Hoffmann 2011). The absence of strong phylogenetic signal among the synthetic chemicals is particularly notable given that most are selective pesticides that were specifically developed to target insects. Instead, non-target organisms, including chordates, appear to be equally as sensitive as insects to many of these chemicals which points to the systemic threat of pesticides to aquatic communities.

There are several factors that might explain the low frequency of strong phylogenetic signal in our results. First, it is possible that the sample sizes of some datasets, like chronic diazinon (*n* = 10 species) and acute 4-nitrophenol (*n* = 14 species), were simply too small to provide adequate statistical power to estimate λ. Sample sizes might increase if data quality collection parameters were less restrictive, however, this solution is not ideal because of the underlying variability in our toxicity datasets. This variation arises because an organism’s sensitivity to a toxicant may be affected by any number of environmental (temperature, pH, salinity, etc.), biological (size, sex, life stage, etc.) or chemical (mode of action, structure, solubility, etc.) factors. We aimed to reduce such variation and its effects by setting rigid requirements for certain variables (CAS number, exposure media, test duration) during data collection, controlling others (life stage, temperature and pH) in the analyses and by using geometric means rather than arithmetic. However, because of the sporadic availability of experimental information in ECOTOX, we left variables such as chemical purity, dissolved oxygen content and test organism sex unrestricted to ensure there were enough data to perform the planned analyses. As such, we expect that some underlying variation remains in our data and may have obscured the phylogenetic signal in some instances.

Additionally, there may be statistical noise in the chronic toxicity data that does not exist in the acute data because of the uncertainty regarding the no observed effect concentration (NOEC). Generated from *post hoc* tests after performing an ANOVA, the NOEC represents the highest concentration of a chemical that does not induce a response that differs significantly from the control, and the biological endpoint used to determine a NOEC can vary. In general, the NOEC is considered to be a relatively poor indicator of safe chemical concentrations (Crane and Newman 2000), and as a result, there has been a strong push in ecotoxicology to utilize alternative measures of chronic toxicity (Warne and van Dam 2008; Laskowski 1995; Kooijman 1996). However, opportunities to work with other types of data are minimal given that chronic data are severely limited or even nonexistent for many chemicals (de Zwart 2001). The paradox created by this shortage and the requirement of existing data for typical CSE on means we are unlikely to be able to comprehensively fill gaps in the chronic toxicity database using this approach. Instead, methods that extrapolate chronic values using acute data (Duboudin, Ciffroy, and Magaud 2004; Hiki and Iwasaki 2020) appear better suited to this challenge. The gaps in the acute toxicity database, however, can be addressed with phylogenetic methods.

The value of phylogenetic approaches to CSE arises from the concept that species data cannot be considered independent observations (Felsenstein 1985). All species are related within a hierarchical phylogenetic tree, so similar phenotype values among species could be a product of limited divergence from a shared common ancestor or convergent evolution (Stone, Nee, and Felsenstein 2011). Most standard statistical tests assume datapoints to be independent, meaning that using such methods to analyze a dataset heavily influenced by evolutionary history could lead to spurious results. Phylogenetically-informed methods avoid this issue by explicitly accounting for phylogenetic structure in a dataset. In this context, the best-studied method of phylogenetic CSE is the phylogenetic eigenvector map (PEM: Guénard, Legendre, and Peres-Neto 2013; Guénard et al. 2014). A PEM is a set of eigenfunctions derived from a phylogeny that describe the magnitude of various possible phylogenetic patterns (i.e. phylogenetic signal values) in a dataset. A subset of these eigenfunctions are then utilized as the independent variables in a regression model that generates estimates of trait values for species on the phylogeny that lack data. PEMs have also been combined with descriptors of chemical properties in a bilinear modelling approach that can predict the toxicity of multiple chemicals to many species (Guénard et al. 2014), which represents a substantial expansion of conventional modelling efforts.

Approaching toxicity data with an evolutionary perspective may benefit pollution management efforts beyond cross-species extrapolation. Here, we identified patterns of sensitivity in the datasets that exhibited strong phylogenetic signal that correspond with descriptions from the literature of how these chemicals affect different taxa. For example, neurotoxins like guthion, an organophosphate insecticide, are typically considered more toxic to invertebrates than vertebrates (Legradi et al. 2018), which is evident in our results (Fig. 3). Similarly, freshwater mussels have been found to be more tolerant of chlorine than other aquatic species (Valenti et al. 2006) which matches with our evaluation of the molluscs in our acute toxicity dataset for chlorine (Fig. 1). These similarities suggest that phylogenetic assessments of toxicity data can provide reliable insights into patterns of sensitivity which, in a regulatory context, could aid in water quality criteria development. For instance, by referring to our guthion plot (Fig. 3), it becomes clear that amphipods are some of the most acutely sensitive taxa to guthion. A toxicity dataset for guthion water quality criteria could then be specifically assembled to include amphipods rather than relying on the EPA’s taxonomic requirements(Stephan et al. 1985) to ensure their representation. While setting unique taxonomic requirements for every chemical is unrealistic, knowledge of patterns of sensitivity could be used to complement other efforts to modernize water quality criteria.

## Conclusions

Given that understanding of the concentrations of chemicals that adversely affect species is a vital component of the risk assessment process, it is critical that we address the major gaps in our toxicity database. Phylogenetic methods of cross-species extrapolation represent a possible alternative to laboratory testing, provided that chemical sensitivity is strongly influenced by evolutionary relationships. We found that strong phylogenetic signal is apparent at high taxonomic levels in a small subset of chemicals in both acute and chronic data, which was surprising given the specific targets of many pesticides included in this study. For those datasets that do feature strong signal, phylogenetic tools provide a framework with which we can reliably assess patterns in chemical sensitivity and a means of avoiding the statistical pitfalls associated with phylogenetically structured data. Moving forwards, we recommend that future efforts in this field evaluate the phylogenetic signal in their data as a preliminary analytical step to ensure the selection of the appropriate model for cross-species extrapolation.

## Acknowledgments

This work was supported by University of Southern California (USC) Sea Grant award NA18OAR4170075.

## Statements and Declarations

### Author Contributions

A.L.C.: Conceptualization, Methodology, Software, Formal analysis, Investigation, Data curation, Visualization, Writing – Original Draft, Writing – Review & Editing. S.E.: Conceptualization, Funding acquisition, Supervision, Writing – Review & Editing.

### Data Availability

Data pertaining to this manuscript are available from Zenodo at DOI: 10.5281/zenodo.8222812.

