## Supplemental Table 1 and Supplemental Figures 1-3 for "Phylogeny predicts tolerance in aquatic animals for only a minority of chemicals"

**Table S1** Toxicity dataset sample sizes before and after filtering for life stage, temperature and pH information

| **Chemical** | **Exposure** | **Dataset Variation** | | | | | | | |
| --- | --- | --- | --- | --- | --- | --- | --- | --- | --- |
|  |  | **Complete** | | **Subadult** | | **Temperature** | | **pH** | |
|  |  | **Toxicity Values** | **Species** | **Toxicity Values** | **Species** | **Toxicity Values** | **Species** | **Toxicity Values** | **Species** |
| Ammonia | Acute | 94 | 48 | 40 | 18 | 73 | 38 | 73 | 37 |
| Cadmium | Acute | 202 | 63 | 101 | 26 | 192 | 62 | 118 | 43 |
| Cadmium | Chronic | 100 | 10 | 15 | 4 | 86 | 8 | 62 | 4 |
| Chlorine | Acute | 101 | 21 | 24 | 6 | 101 | 21 | 99 | 20 |
| Copper | Acute | 240 | 67 | 103 | 33 | 215 | 65 | 191 | 49 |
| Copper | Chronic | 289 | 26 | 95 | 12 | 183 | 20 | 170 | 20 |
| Mercury | Acute | 73 | 25 | 49 | 10 | 71 | 23 | 35 | 15 |
| Nickel | Acute | 50 | 21 | 25 | 7 | 36 | 18 | 28 | 13 |
| Phenol | Acute | 167 | 55 | 39 | 16 | 148 | 55 | 134 | 44 |
| Toluene | Acute | 70 | 21 | 33 | 10 | 70 | 21 | 55 | 13 |
| Zinc | Acute | 148 | 49 | 69 | 15 | 133 | 47 | 84 | 32 |
| 4-Nitrophenol | Acute | 42 | 14 | 20 | 5 | 42 | 14 | 25 | 10 |
| Atrazine | Acute | 109 | 53 | 44 | 25 | 109 | 53 | 79 | 41 |
| Atrazine | Chronic | 350 | 43 | 132 | 25 | 309 | 37 | 207 | 24 |
| Chlorpyrifos | Acute | 303 | 100 | 101 | 49 | 300 | 98 | 172 | 75 |
| Chlorpyrifos | Chronic | 210 | 36 | 53 | 16 | 154 | 27 | 110 | 18 |
| DDT | Acute | 368 | 99 | 62 | 30 | 368 | 99 | 213 | 76 |
| Diazinon | Acute | 203 | 83 | 76 | 42 | 196 | 81 | 130 | 59 |
| Diazinon | Chronic | 126 | 10 | 9 | 5 | 80 | 8 | 68 | 6 |
| Dieldrin | Acute | 173 | 65 | 66 | 28 | 157 | 60 | 139 | 57 |
| Endosulfan | Acute | 363 | 89 | 59 | 31 | 311 | 84 | 232 | 69 |
| Endrin | Acute | 202 | 66 | 46 | 25 | 201 | 65 | 143 | 57 |
| Glyphosate | Acute | 98 | 30 | 38 | 16 | 95 | 29 | 62 | 18 |
| Glyphosate | Chronic | 101 | 11 | 39 | 8 | 96 | 9 | 50 | 4 |
| Guthion | Acute | 245 | 54 | 54 | 23 | 242 | 53 | 197 | 49 |
| Lindane | Acute | 220 | 93 | 46 | 28 | 211 | 91 | 181 | 84 |
| Malathion | Acute | 343 | 132 | 101 | 58 | 340 | 131 | 278 | 110 |
| Malathion | Chronic | 137 | 12 | 40 | 8 | 120 | 12 | 102 | 6 |
| Parathion | Acute | 146 | 50 | 58 | 25 | 146 | 50 | 86 | 37 |
| PCP | Acute | 356 | 99 | 128 | 44 | 327 | 88 | 309 | 81 |
| PCP | Chronic | 102 | 23 | 65 | 17 | 81 | 19 | 64 | 20 |
| TBTO | Acute | 69 | 33 | 24 | 12 | 66 | 33 | 53 | 29 |

Abbreviations: DDT = 1,1′-(2,2,2-Trichloroethane-1,1-diyl)bis(4-chlorobenzene), PCP = Pentachlorophenol, TBTO = Tributyltin oxide

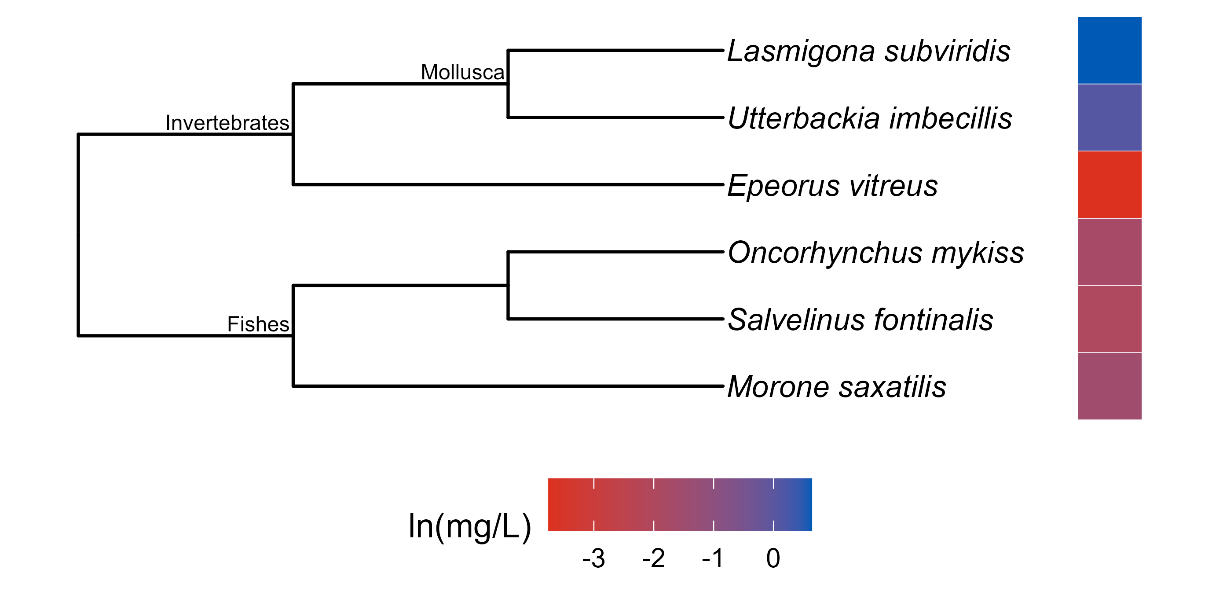

**Fig. S1** Phylogenetic tree and toxicity data heatmap for the acute chlorine subadult dataset (λ = 0.68). The colored bar next to each species represents its relative sensitivity to the chemical. A red bar indicates a high degree of sensitivity (i.e. small amount of chemical causes toxic effect), while a blue bar indicates low sensitivity (i.e. large amount of chemical causes toxic effect).

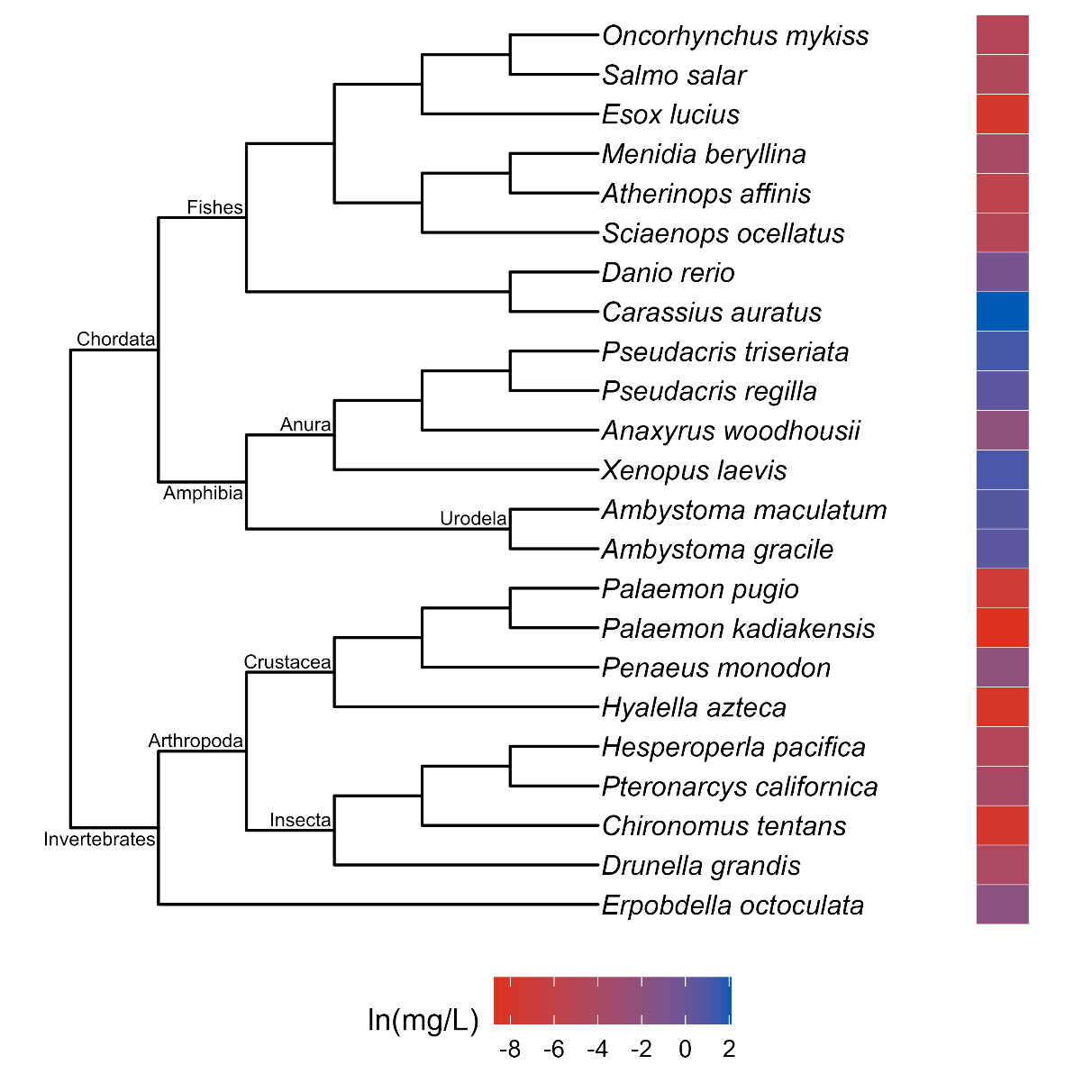

**Fig. S2** Phylogenetic tree and toxicity data heatmap for the acute guthion subadult dataset (λ = 0.69). The colored bar next to each species represents its relative sensitivity to the chemical. A red bar indicates a high degree of sensitivity (i.e. small amount of chemical causes toxic effect), while a blue bar indicates low sensitivity (i.e. large amount of chemical causes toxic effect).

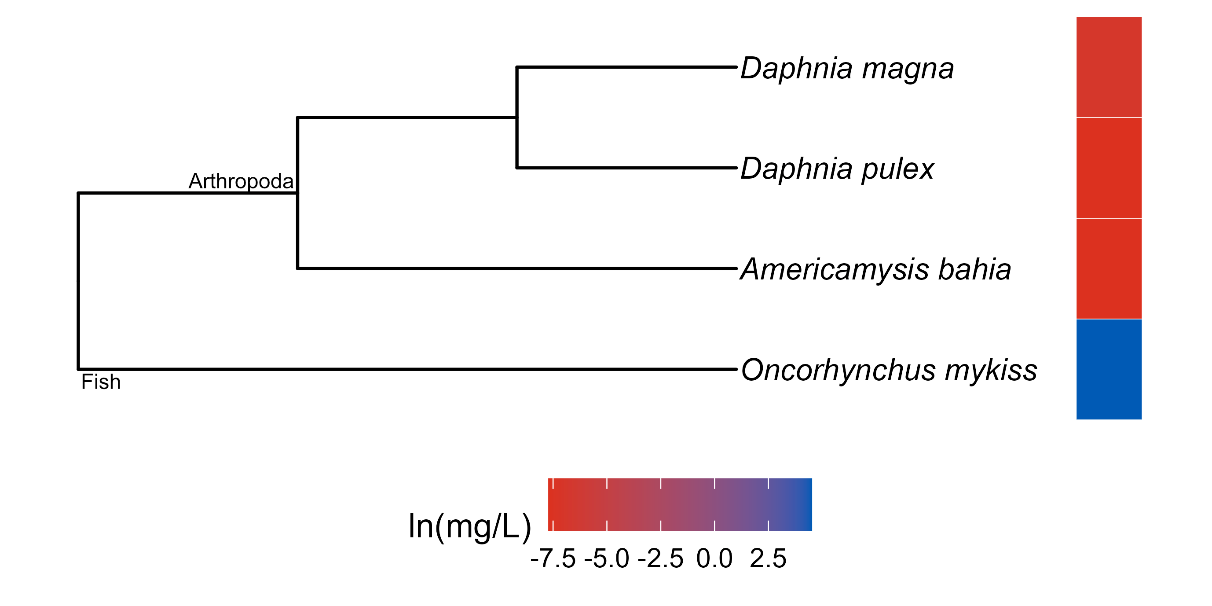

**Fig. S3** Phylogenetic tree and toxicity data heatmap for the chronic cadmium subadult dataset (λ = 1.0). The colored bar next to each species represents its relative sensitivity to the chemical. A red bar indicates a high degree of sensitivity (i.e. small amount of chemical causes toxic effect), while a blue bar indicates low sensitivity (i.e. large amount of chemical causes toxic effect).
